# Simulation-Based Inference at the Theoretical Limit: Fast, Accurate Microstructural MRI with Minimal diffusion MRI Data

**DOI:** 10.1101/2024.11.11.622925

**Authors:** Maximilian F. Eggl, Silvia De Santis

## Abstract

Diffusion-weighted magnetic resonance imaging (dMRI) is an essential tool in neuroscience, providing non-invasive insights into brain microstructure. However, obtaining accurate maps demands long acquisitions, due to oversampling of the parameter space. Leveraging simulation-based inference (SBI) with neural networks, we directly approximate posterior distributions of diffusion parameters from experimental measurements without requiring real-data training. SBI achieves accurate parameter estimation using up to 90% fewer acquisitions and outperforms standard nonlinear least squares under noisy, sparse sampling. We demonstrate improvements across diffusion tensor imaging, diffusion kurtosis imaging, and biophysical models estimating axonal density and calibre. Validation on simulated and real datasets from healthy and pathological brains shows SBI’s robustness and generalizability. This approach can expand dMRI access, e.g. for paediatric and other time-sensitive patients, enable advanced microstructure-sensitive protocols, and rescue legacy data with suboptimal quality. By reducing scan times and preserving privacy, SBI-integrated dMRI workflows promise faster, more comfortable exams, MRI-based virtual tissue biopsy, and have potential to substantially impact radiology workflows.

## 1 Introduction

By sensitizing magnetic resonance imaging (MRI) signals to water’s random motion, diffusion-weighted MRI (dMRI) has become a pivotal neuroimaging tool, providing unique insights into tissue microstructure in vivo and noninvasively [1, 2]. When combined with mathematical representations based on tensors like diffusion tensor imaging (DTI) [3] or diffusion kurtosis imaging (DKI) [4], dMRI has significantly impacted clinical applications [5, 6]. More recently, advances aiming at dissecting the MRI signal coming from distinct brain tissue compartments has led to the development of biophysical representations, which notably allow for direct measurement of fine cell properties like soma size, neurite morphology, permeability and axon radius [7–13]. These insights, thus, provide a non-invasive way of deriving meaningful biomarkers of health and disease, effectively positioning dMRI as a possible tool in achieving virtual tissue biopsy. However, noisy acquisitions substantially affect these methods, compromising the accuracy and reliability of fitted diffusion parameters [14, 15]. While denoising techniques exist (e.g., [16–19]), fitting methods still demand extensive oversampling— lengthening scan times by up to 10 times the theoretical minimum or more, excluding some patients, and precluding richer multimodal protocols [20–22]. In fact, the vast majority of toolboxes and software used to extract parameter maps from dMRI data utilize least square approaches, statistical frameworks which determine the optimal parameters of a model by minimizing the sum of the squared differences between observed and predicted values. However, these methods, including non-linear least squares (NLLS) and weighted least squares (WLS) [3, 19, 23], besides requiring massive oversampling, are additionally sensitive to initial parameter guesses and may not fully capture uncertainties in noisy, low signal-to-noise ratio (SNR) conditions [14]. More modern approaches like Rician-unbiased or variational Bayesian [24, 25] have been introduced to mitigate these issues. However, their high computational cost, reliance on precise noise-variance estimates under a strict Rician model, and tendency toward overconfident posteriors in low-SNR regions have so far limited their practical adoption. Due to these limitations, even basic diffusion models remain underused in clinical practice, and more insightful methods rarely reach clinical researchers. A huge opportunity to translate recent MRI advances into clinical practice is lost in the process.

Simulation-based inference (SBI) sidesteps these issues by using simulated acquisitions to learn posterior distributions of diffusion parameters [26–29]. Neural posterior estimation (NPE) [30, 31] trains a deep network on simulated data, yielding robust fits even in low-SNR regimes. NPE has been successfully applied in fields like cosmology [32], neuroscience [33, 34], and imaging neuroscience [35–38]. Nonetheless, in the context of dMRI this framework has not been used previously to explore the idea of achieving accurate parameter maps with the minimal amount of theoretically required information. Importantly, SBI relies on simulations, contrasting with other deep learning approaches that require training on real data [39, 40], making it data- and privacy-friendly.

Since the diffusion-based representations we study commonly have straightforward simulation paradigms, SBI, and particularly NPE, is a natural fit for enhancing fitting accuracy. This work demonstrates how applying SBI to different tensor and non-tensor based dMRI representations allows direct fitting of the models and extraction of accurate parameter maps with minimal or significantly reduced acquisitions. Here, we focus on four different established models: DTI, DKI, Composite Hindered And Restricted ModEL of Diffusion (CHARMED) and AxCaliber, but the approach is inherently flexible and readily extendable to emerging or future models. By validating the capabilities of SBI and dMRI using simulated and real scans from healthy and pathological datasets and across different sampling schemes [41, 42], we believe this approach can substantially impact radiology in research and clinical settings, providing a framework to extract the maximum amount of microstructural information with minimum acquisition time and revolutionize accessibility to both basic and advanced dMRI.

## 2 Methods

### 2.1 Tensor-based frameworks

In DTI, diffusion is modelled as a Gaussian process using the diffusion tensor *D*, a 3 × 3 symmetric positive-definite matrix representing diffusion coefficients in three-dimensional space. The signal attenuation is given by:

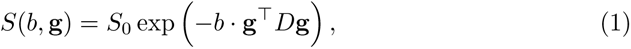

where *S*(*b,* **g**) is the measured signal with diffusion weighting factor *b* and gradient orientation **g**, and *S*_0_ is the non-diffusion-weighted signal. This framework allows derivation of metrics related to the diffusion tensor like mean diffusivity (MD) and fractional anisotropy (FA) (Fig. 1a), widely employed for diagnostic purposes (e.g. [43]; review: [44]). However, the Gaussian assumption underlying DTI limits its sensitivity to more complex diffusion phenomena.

**Fig. 1:**
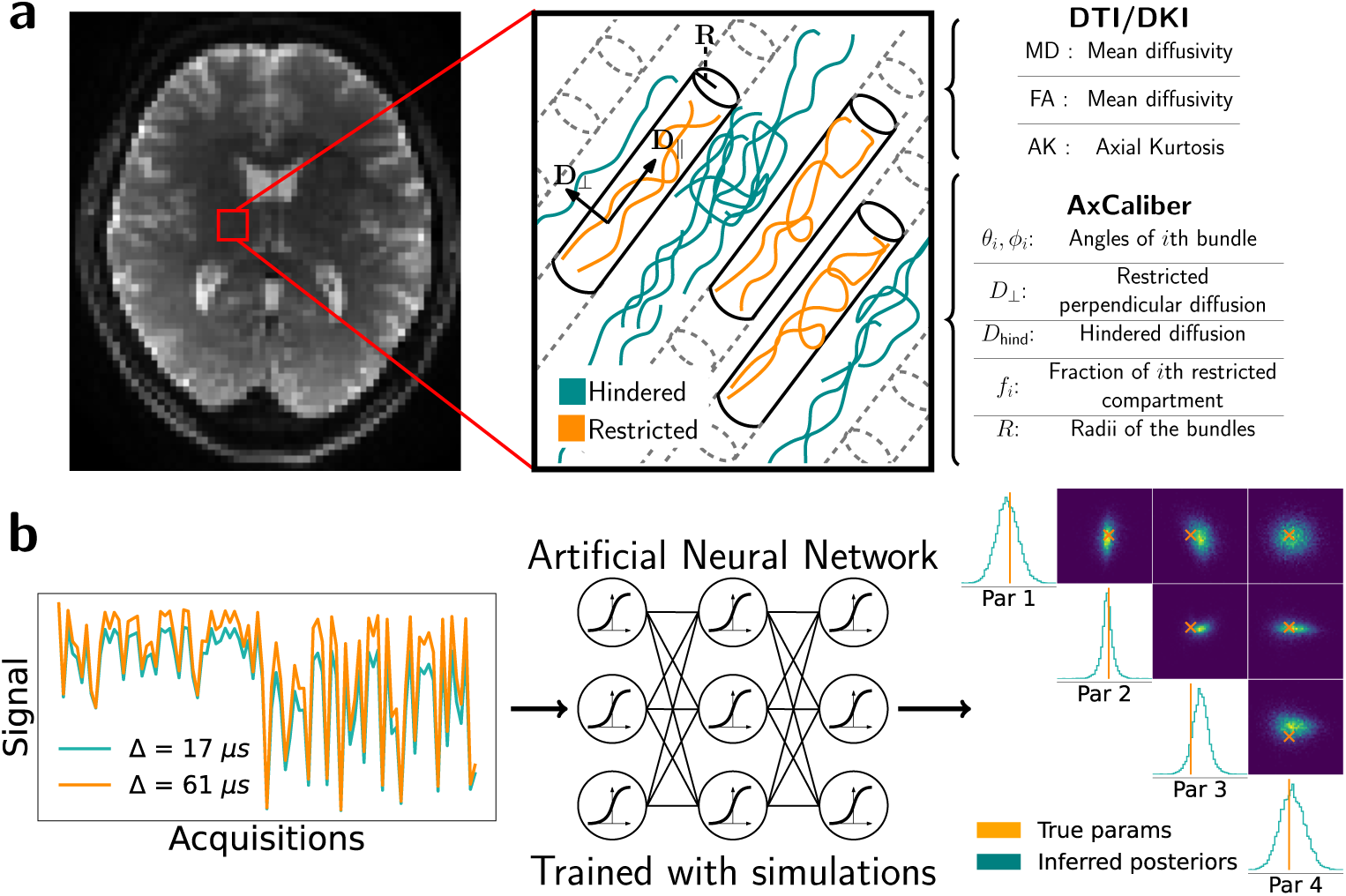
Overview of the dMRI data processing and analysis pipeline. **a)** dMRI data from the brain captures the diffusion of water molecules as they interact with the underlying brain microstructure. This data can be analysed with various models, for example, mathematical signal representations such as DTI or DKI or more advanced multi-compartment models such as CHARMED and AxCaliber. These models then provide interpretable biomarkers which give insight into the underlying health of the brain parenchyma. **b)** When sampling different gradient strengths, gradient directions, and diffusion times, the signal acquires a fingerprint of the underlying microstructure. By using a simulation-based inference approach, this signal is provided to a network pre-trained on simulated signals generated by a model (e.g., DTI). This network then generates the posterior distribution over the model parameters given the measured diffusion signal.

To overcome this limitation, DKI extends the DTI model by incorporating non-Gaussian effects, capturing the kurtosis of water diffusion. The signal attenuation in DKI is modelled as:

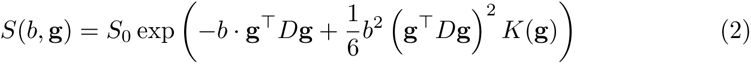

where *K*(**g**) is the kurtosis coefficient in orientation **g** [4]. DKI captures more complex diffusion dynamics, providing metrics like mean kurtosis (MK) for deeper microstructural insights. Other DKI metrics include axial kurtosis (AK), radial kurtosis (RK), the mean kurtosis tensor (MKT), and kurtosis fractional anisotropy (KFA). MKT and KFA, though less common, specifically assess the accuracy of kurtosis tensor estimates [45, 46].

### 2.2 The AxCaliber model

Here, we briefly introduce the AxCaliber framework, an extension of the CHARMED formulation (for full derivations, see [12, 47]). AxCaliber models the signal as arising from two compartments: a hindered component (denoted as (*·*)*_h_*) and a restricted component (denoted (*·*)*_r_*), as illustrated in Fig. 1a. For the hindered component, we use the standard DTI framework with *D_h_* as the diffusion tensor. The restricted compartment, representing white matter microstructure, is modelled as radially symmetric cylindrical structures (i.e., no radial diffusion). Its signal is decomposed into parallel and perpendicular components. Assuming the cylinder is long relative to the diffusion distance, the parallel component simplifies to a simple 1-D exponential decay with *D*_‖_ as the scalar diffusion coefficient. The perpendicular component, influenced by the cylinder’s radius, *R*, is significantly more complex, and the full expression is omitted here but can be found in [48]. A key aspect of AxCaliber is recognizing that axon bundles feature a distribution of radii. Originally a gamma distribution was used, but difficulties in fitting both shape and scale motivated our use of a Poisson model [49]. This model requires fitting only the mean radius *λ*, simplifying interpretation.

Thus, the complete expression of the diffusion signal becomes

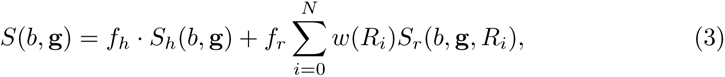

where *f_h_, f_r_* are the fractions of the hindered and restricted components, *S_h_, S_r_* are the diffusion signals of those compartments and *w*(*R_i_*) are the normalized weights from the Poisson distribution. Importantly, the AxCaliber formulation posits that varying the diffusion time Δ (with other parameters adjusted to keep the same b-value) yields different signals, as shown in Fig. 1b. Intriguingly, the simplified Poisson formulation requires fitting only a single parameter (similar to the ActiveAx implementation [50], which only fits the mean axon diameter), theoretically enabling axon diameter estimation from a single diffusion time.

### 2.3 Simulation-based inference fitting framework

In this work, we employed NPE [31, 51], an approach in the family of SBI-methods to fit the parameters of a given diffusion model. NPE leverages normalizing flows [52], flexible probabilistic models constructed by applying invertible transformations to simple distributions, such as Gaussians. By utilising neural networks to learn the normalizing-flow transformations, NPE can approximate the posterior over parameters even when the likelihood is intractable.

Fig. 1b provides a schematic overview of the NPE procedure. Briefly, during training, candidate parameters are sampled from a prior distribution, and synthetic signals are generated by applying these parameters to our forward DTI, DKI or AxCaliber models. The NPE network is optimized by minimizing the negative log probability of the sampled parameters under its predicted posterior, conditioned on each synthetic signal. After convergence, we input experimental data into the trained network to obtain the posterior distribution of model parameters. Since the network outputs a full posterior distribution given the input data, it naturally provides a measure of uncertainty that reflects the confidence in our parameter estimates. To obtain a single parameter estimate from the posterior, several approaches can be adopted, such as taking the mean (when the posterior is well-behaved), the mode (histogram mode), or employing more advanced techniques like Maximum a Posteriori (MAP) estimation [53]. In this work, we used the posterior mean, computed efficiently by averaging a large batch of parallel samples, eliminating additional voxel-wise optimization and scaling easily to whole-slice inference.

### 2.4 In silico data

As SBI relies on simulated data for training, realistic simulations are crucial for accurate results [54, 55]. Therefore, for the three investigated models, the priors used to sample the relevant parameters were designed to mimic features of real human data [28, 56].

For DTI simulations, parameters were sampled from uniform distributions and adjusted to ensure the diffusion tensor was positive definite and symmetric. It was found that the uniform distribution for the diagonal entries needed to be roughly twice as wide as that for the off-diagonal entries. With this approach, we were able to effectively replicate the HCP data features without prior knowledge (see Supplementary Fig. S1).

DKI simulations required a more sophisticated approach due to the non-linear relationship between DTI and DKI terms and challenges in defining suitable priors. Generating stable signals was crucial because DKI includes a positive *b*^2^ term, which can lead to unstable simulations if *D* and *K* are not carefully chosen. To address this, prior distributions were obtained from a single subject of the HCP dataset (excluded from further analysis) to generate diffusion and kurtosis tensors that reflect real-world non-linear relationships while remaining biologically plausible. More precisely, where appropriate, we fit log-normal or normal distributions to each of the entries of the diffusion and kurtosis tensors (see Supplementary Figs. S2b-d). Approximately 15% of tensors yielded unstable signals due to mathematically invalid tensor–kurtosis combinations (not biological variability) and were resampled.

Finally, the AxCaliber framework models a combination of diffusion tensors (for the hindered component) and biologically driven parameters. Diffusion tensors followed the same procedure as above, while priors for biological parameters were uniform distributions informed by literature (see for example axon size [57, 58]).

Simulated signals were generated using a Monte Carlo approach and corrupted with Rician noise at varying signal-to-noise ratios [59]. Simulations were split into training and testing sets; neural networks were trained on the training set to estimate model parameters, and the testing set was used to compare fitted parameters of both SBI and NLLS against the available ground truth. All simulations and tensor-specific metric calculations were performed using Dipy functions [60], while we implemented an in-house Python implementation of the AxCaliber framework.

We generated 300,000 training samples for each model. Unless stated otherwise, in silico inference results were generated using 500 simulated signals. DKI priors were fitted using `scipy.stats`, and NPE was performed with the `sbi-toolbox` [29].

### 2.5 Human dMRI data

To validate the performance of the SBI framework for fitting DTI and DKI parameters using real-world data, we selected 32 individuals (17 females and 15 males, ages 20-55) from the Human Connectome Project (HCP) database. Initial pre-processing was performed by the HCP [Minimal Preprocessing Pipeline 61], while the study-specific pre-processing consisted of downsampling the images from 1.5 mm/voxel to 2.5 mm/voxel. This downsampling was performed for the sake of computational efficiency, but all results in this work also apply to the higher 1.5 mm/voxel resolution (see supplemental fig. S3). The datasets consisted of two sets of b-values, *b* = 1000 and *b* = 3000, with 69 gradient acquisitions per b-shell.

To validate the DKI fits on a different dataset as well as study the AxCaliber model, which requires multiple diffusion times, we selected 8 additional individuals (6 females and 2 male, ages 26-55, 5 with MS and 3 healthy subjects) from the dataset of [42]. This data, available through the public repository DIGITAL.CSIC and pre-processed with the same HCP pipeline, consisted of two sets of b-values, *b* = 2000 and *b* = 4000, with 30 and 59 gradient acquisitions, respectively. Diffusion times (Δ) were utilised in the AxCaliber setting and were set to 7, 35, and 61 *ms*. Images were also downsampled to 2.5 mm/voxel as above.

### 2.6 Number of acquisitions used

A key focus of this work is evaluating how SBI and NLLS performance varies with the number of acquisitions used under DTI, DKI, and AxCaliber models. To this end, we defined the following acquisition subsets:

- **DTI full set**: 69 acquisitions (5 *B*_0_, and 64 with *b* = 1000)
- **DTI medium set**: 20 acquisitions (1 *B*_0_, and 19 with *b* = 1000) which, according to [21], represents a realistic and near-optimal set of 20 orientations sufficient for the DTI framework.
- **DTI minimum set**: 7 acquisitions (1 *B*_0_, and 6 with *b* = 1000, theoretical
- minimum)
- **DKI full set**: 138 acquisitions (10 *B*_0_, 64 with *b* = 1000, 64 with *b* = 3000)
- **DKI medium set**: 35 acquisitions (1 *B*_0_, 6 with *b* = 1000, 28 with *b* = 3000)
- **DKI minimum set**: 22 acquisitions (1 *B*_0_, 6 with *b* = 1000, 15 with *b* = 3000)
- **AxCaliber full set**: 271 acquisitions (1 *B*_0_, 30 with *b* = 2000, 60 with *b* = 4000, repeated for three diffusion times)
- **AxCaliber reduced set**: 19 acquisitions (1 *B*_0_, 2 with *b* = 2000, 3 with *b* = 4000, repeated for three diffusion times)

Gradient orientations in all simulated acquisition sets were selected using electrostatic repulsion principles [20] to ensure optimal angular coverage, primarily to support NLLS. Notably, SBI remained robust even under suboptimal orientation and b-value configurations, underscoring its suitability in data-limited settings (see Supplemental Fig. S4).

### 2.7 Metrics to evaluate SBI and NLLS performance

In our in-silico simulations, access to the ground truth allows direct comparison between estimated and true metrics. We quantified error as the absolute difference between true and estimated metrics.

For human data, lacking ground truth, we compared results from minimal and medium acquisition sets against the full set. By working with images rather than isolated signals, we leveraged structural information to assess performance. As a proxy for accuracy, we used the Structural Similarity Index (SSIM) [62], which measures similarity based on luminance, contrast, and structure. SSIM values closer to 1 indicate higher similarity; we set a minimum desired performance threshold of 66% based on clinical interpretability criteria, as preliminary analyses showed that SSIM values above this level preserved structural features necessary for clinical interpretation. Such features include lesion boundaries and tissue interfaces, ensuring minimal perceptible differences for clinicians. However, we note that most of our SBI results reach a much higher benchmark (often *>* 90%), meaning that the results become virtually indistinguishable from the ground truth versions.

To assess precision, we calculated the standard deviation of the fitted metric within white matter. We considered performance acceptable when the interquartile range of minimum or reduced acquisition fits matched that of full acquisitions.

## 3 Results

### 3.1 Validating SBI in DTI and DKI

#### 3.1.1 In-silico analysis

For DTI, we began by generating a single signal and adding noise at varying SNR levels (Fig.2a). Estimating MD and FA from the 200 noisy versions, we found that NLLS was accurate at SNR *>* 10 but declined at lower SNR, while SBI remained robust down to SNR = 2 with lower variance (Fig.2b).

Across 500 noisy signals, both methods performed similarly at high SNR with full acquisitions (Fig.2c). At lower SNR, SBI’s errors remained lower. With minimum acquisitions (Fig.2d), MD estimates remained stable, but FA from NLLS degraded substantially, while SBI retained better accuracy. Supplemental Fig. S5 confirms that SBI reconstructs the true signal more faithfully, while NLLS tends to overfit the noise (higher correlation with the noisy signal than SBI in a iv) and vi) of the supplemental figure).

Using simulated DKI data based on HCP characteristics (Supplementary Fig. S6), we first compared SBI and NLLS using the full set of acquisitions. With a single signal at SNR = 20, both methods reconstructed the signal effectively (Fig. 2e). Across 500 noisy signals, SBI outperformed NLLS at all SNR levels, maintaining higher robustness (Fig. 2f).

**Fig. 2:**
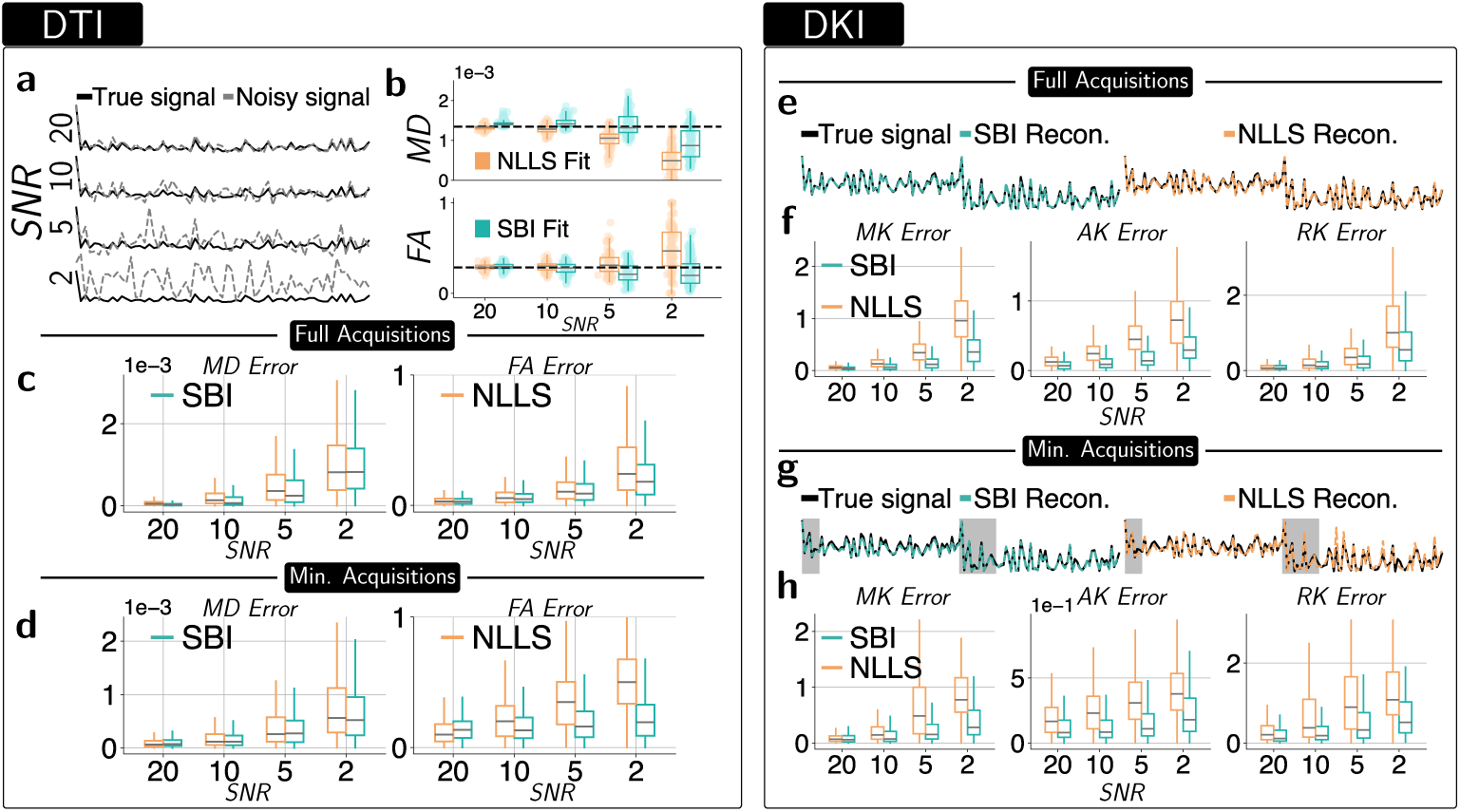
SBI applied to simulated data of DTI and DKI models shows more accurate performance than standard NLLS. **a)** dMRI data is affected by varying levels of noise, which impacts signal clarity. The effect of different signal-to-noise ratios (SNR = 2–20; dashed gray) is shown in comparison to the true signal (black). **b)** Accuracy of estimates of DTI metrics (MD - top and FA - bottom) obtained using the SBI (teal) and NLLS (orange) approach. The black dashed line indicates the true parameter values while the box plots represent median values, with boxes showing IQR and whiskers extending to 1.5 times the IQR. **c-d)** Using either the full set of acquisitions (c) or the minimum (d), both NLLS and SBI are fit to 500 simulated signals with the absolute difference from the true values in MD and FA being shown. Box plots as in (b). **e)** Under the DKI framework, SBI and NLLS are used to infer model parameters and reconstruct an example DKI signal from the full acquisition set (SNR = 20). **f)** To evaluate performance across a range of simulated signals, 500 example DKI signals were generated and subjected to four SNRs (20, 10, 5, and 2). The resulting parameter fits were compared to the true parameters. Box plots as in (b). **g-h)** Repeating the analysis in panels e–f with the minimum DKI set of acquisitions.

Next, we used the minimum set of acquisitions for the DKI inference. In reconstructing the signal (Fig. 2g), SBI achieved more accurate results despite limited data, especially at higher b-values where NLLS deviated noticeably from the true signal. When comparing across 500 signals (Fig. 2h), in general both methods performed worse with minimal data than with full measurements due to reduced information. Interestingly, as SNR decreased, NLLS performance sometimes improved with fewer measurements—consistent with DKI’s known sensitivity to noise, where additional noisy data can degrade fits. In contrast, SBI maintained stable performance across SNR levels and acquisition sizes, effectively extracting reliable information even from minimal data.

#### 3.1.2 In-vivo analysis

To mimic clinical application, we examined MD and FA maps from a medial axial slice of one HCP subject, comparing SBI and NLLS (Figs. 3a,c). Both methods yielded similar structures, though NLLS had lower MD values in the lateral ventricles. Based on trends from in-silico data — where NLLS tended to underestimate MD — we argue that SBI likely provides a more accurate reflection of the true tensor metrics. For FA, both methods yielded similar results, with differences driven primarily by noise.

**Fig. 3:**
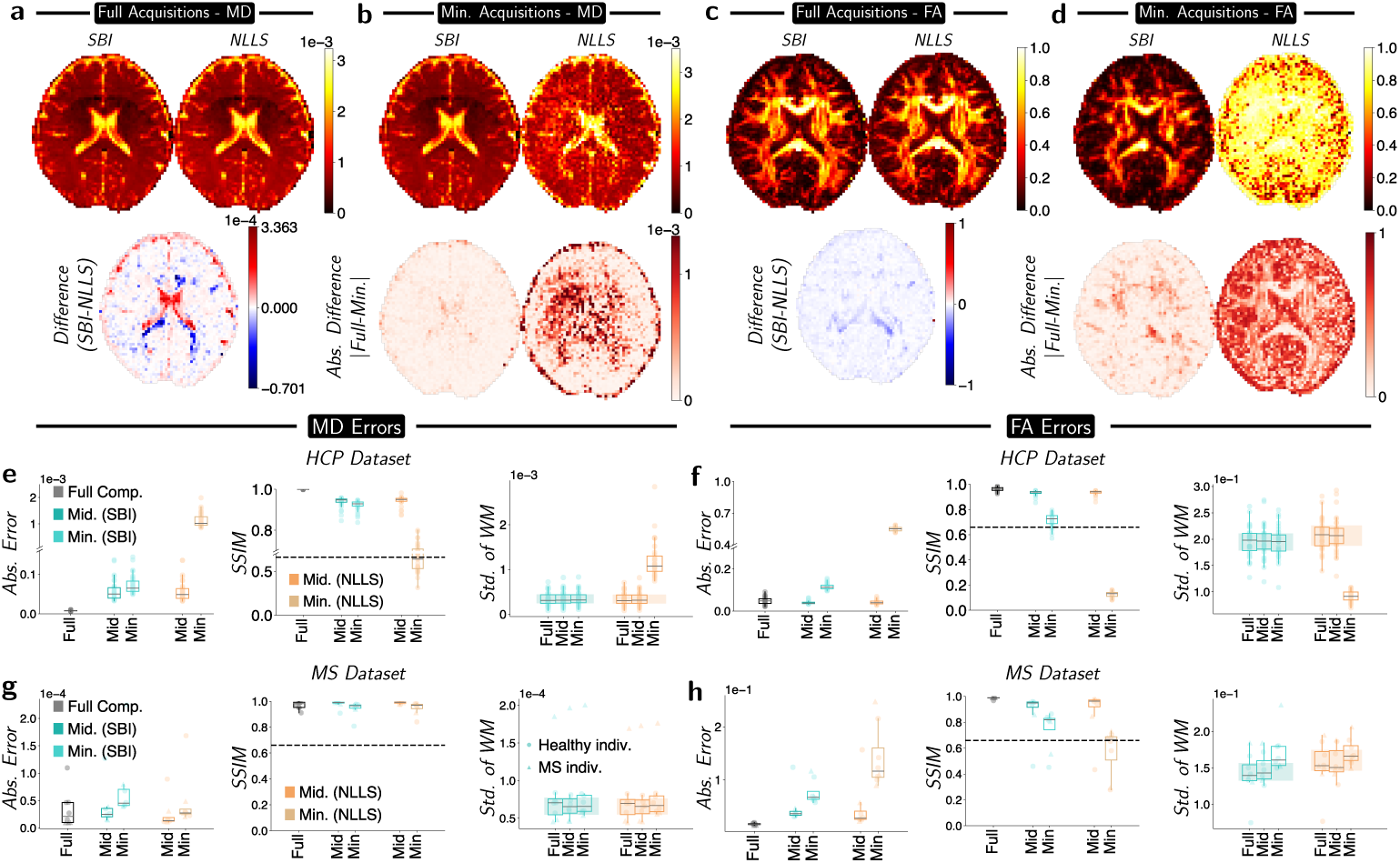
Even with minimal information SBI is able to get meaningful estimates on real-world data where NLLS fails. **a-b)** Comparison of MD values obtained with SBI and NLLS using the full and minimal acquisition sets, respectively (differences below). **c-d)** Similar to (a–b), but showing FA instead of MD. **e-f)** MD and FA were fit with SBI (single network, teal) and NLLS (orange) on medial-axial slices from 31 HCP subjects, using full, medium, and minimal acquisition sets. We report absolute errors in MD and FA, SSIM, and the standard deviation of each metric in white matter. Given a lack of an objective ground truth, the full NLLS and full SBI results are used for their respective subsets. A comparison between the full SBI and full NLLS is shown in the dark gray boxplot. Boxplots represent the median values while the boxes represent the IQR and the whiskers 1.5 the IQR. The dashed black line in the SSIM plots refers to our acceptable minimum accuracy. The shaded areas in the standard deviation plots refer to the interquartile range of the full-set result. **g-h)** The same analysis as in e, f but performed for 8 individuals (3 control and 5 with MS) of the [42] dataset.

With minimal measurements (Figs. 3b,d), SBI’s advantage becomes clear. SBI’s MD map closely matches the full version (SSIM = 0.97), whereas NLLS appears noisier (SSIM = 0.59). Even the more complex FA metric is effectively reproduced by SBI, whereas NLLS loses structural information.

We then trained a generalist network to simultaneously evaluate 31 HCP subjects. Without ground truth, we compared each subset against its own full fit. The results of the absolute error, SSIM, and white matter standard deviation of the metrics (when compared against the respective full fit) are shown in Fig. 3e and f. For SBI, the accuracy of both MD and FA remains well above the 66% minimum accuracy we set (with the minimum SBI MD only losing approximately 5.6% SSIM accuracy with less than 10% of the available information) and far outperforms the minimum NLLS results (65% average SSIM MD accuracy loss). SBI maintained consistent precision in white matter; NLLS deviated from the full acquisition IQR under minimal conditions and showed artificially low FA variability in the minimum acquisition setting due to producing uniform (and incorrect) values. We note that a similar picture appears in the [42] dataset that includes both healthy and pathological individuals (Fig. 3g-h). Here both NLLS and SBI perform well for the MD metric, even under the sparsest conditions. However, NLLS is unable to obtain accurate FA measurements for this case, which SBI is still able to do. Importantly, we note that the b-values of that dataset (b=2000) are not optimal for the purposes of DTI. Nonetheless, SBI is able to obtain reasonable parameter estimates even with this sub-optimal paradigm.

To ascertain if applying denoising (in this case MP-PCA[63] to the dataset before fitting NLLS would improve its performance, we took an individual from the HCP dataset and added noise at varying strengths (SNR = 50 to 10) and then denoised the image and fit NLLS - on the other hand SBI was directly applied to the corrupted images. The results of this are seen in supplemental figure S7. We note that SBI is able to perform the fit on the noisy results where NLLS still fails.

Next, we repeated this analysis for the DKI model. With the full acquisition set, both SBI and NLLS captured similar structural features across kurtosis metrics (Fig. 4a-b). Importantly, the kurtosis-specific metrics, such as MKT and KFA, were well resolved by both methods (see Fig. S8a). However, SBI showed greater stability, while NLLS produced gaps from unrealistic or negative values, visible as black dots.

**Fig. 4:**
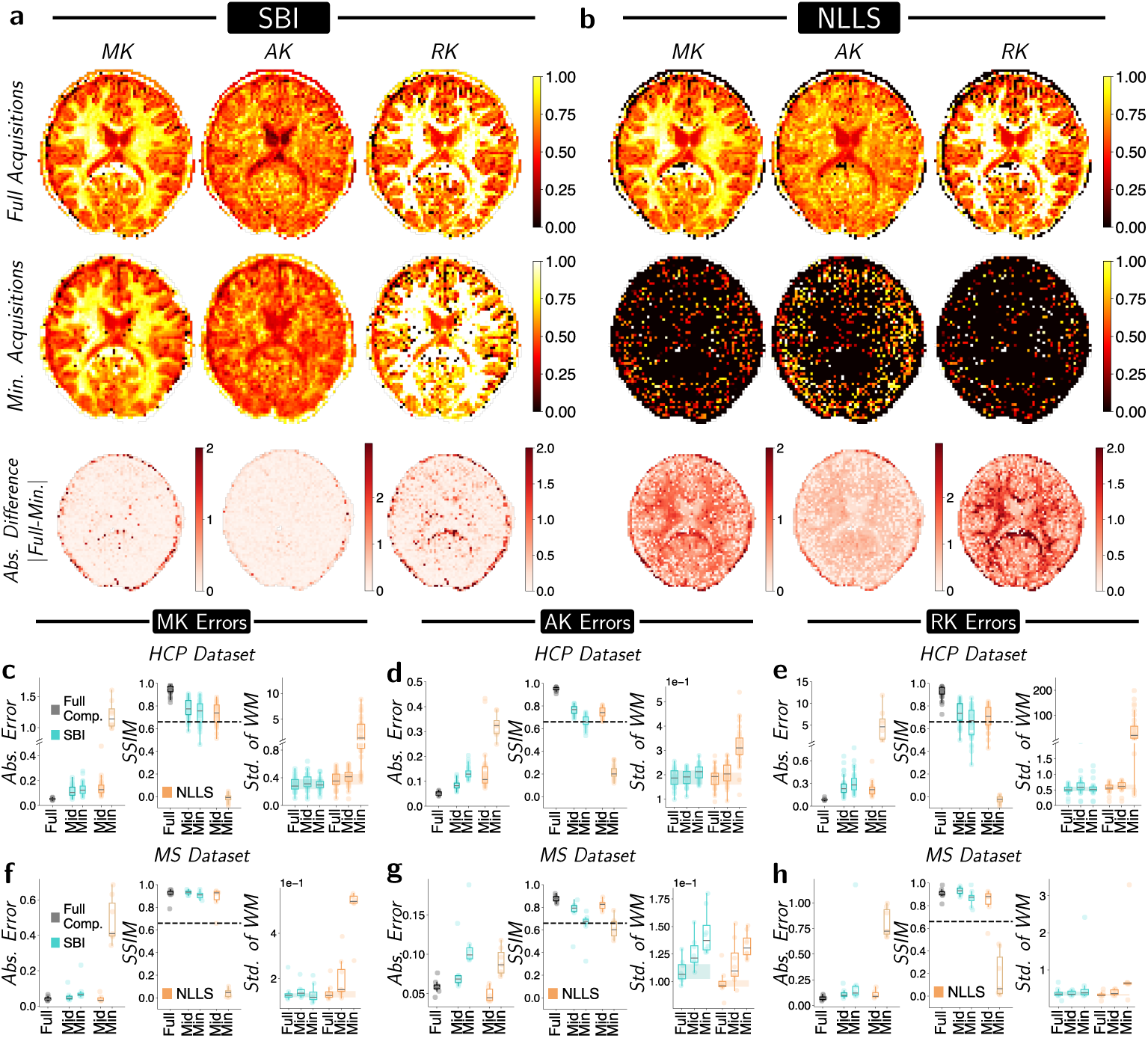
Even with minimum acquisitions, SBI can resolve kurtosis metrics on real-world data where NLLS fails. **a-b)** SBI (a) and NLLS (b) were applied to the axial slice of a held-out individual using full and minimum acquisition sets. This enables qualitative comparison of fitted kurtosis tensor metrics, with black dots marking large negative or invalid values. Differences between full and minimum fits (for both SBI and NLLS) are shown as heat maps (darker red = larger error). **c-e)** SBI (teal) and NLLS (orange) were fit to medial-axial slices from 31 HCP subjects using full, medium, and minimum acquisition sets (via a single network). Panels show absolute error, SSIM, and standard deviation of each metric in white matter, analogous to the DTI analysis. Full-set SBI and NLLS results serve as references for their corresponding reduced-set fits; gray boxplots compare full SBI vs full NLLS for error and SSIM. Boxplots show medians; boxes, IQR; whiskers, 1.5× IQR. Orange denotes NLLS; teal, SBI. Dashed black lines in SSIM plots mark our minimum-acceptable accuracy. Shaded areas in standard deviation plots indicate the IQR of the full acquisition result. **f-h)** The same analysis as in c-e but performed for 8 individuals (3 control and 5 with MS) of the [42] dataset.

Considering the minimum acquisition setting, SBI maintained high-quality reconstructions across DKI metrics. In contrast, NLLS failed under these conditions, producing implausibly negative values. Compared to SBI, NLLS exhibited greater structural discrepancies between full and minimum results, indicating NLLS’s inability to recreate accurate tensor metrics with limited data.

As before, we trained a network capable of fitting DKI to 31 HCP individuals simultaneously, with the resulting absolute error, SSIM, and standard deviations of the metrics in Fig. 4c-e. The increase in absolute error from medium to minimum acquisitions was substantially smaller for SBI than for NLLS. Additionally, SBI achieved higher SSIM scores than its NLLS counterpart. SBI also maintained stable precision, unlike NLLS, which degraded with fewer measurements. To address potential bias from using a single HCP subject for simulation priors, we tested generalizability using a different dataset[42] in Fig. 4f-h. In particular, we applied the same protocol to both healthy individuals and those exhibiting MS. Across the board, SBI was able to get accurate DKI metrics (*>* 0.85 SSIM score for the minimum set for MK and RK) that mirrored the results of the HCP result.

Importantly, due to the non-linearities of the DKI models, we used it to run a more extensive benchmarking of SBI against other fitting strategies, less commonly adopted than NLLS, but potentially more stable at low SNRs, namely weighted least square, Maximum likelihood estimation (MLE) and a variational Bayesian approaches. We compared the performance of all approaches in both simulations and the HCP dataset. In all cases, SBI was able to outperform these methods, both for the full and minimum sets (see Supplemental Fig. S9).

### 3.2 Application of SBI to AxCaliber

We evaluated AxCaliber performance (and by extension, CHARMED) using both in-silico simulations and in-vivo data by first testing the SBI framework on synthetic signals with known ground truth, followed by application to real-world data from the [42] dataset. To start, we generated a set of 500 noisy signals and then fitted AxCaliber to each with SBI and NLLS. The average difference between the inferred angles of the restricted component and the true angles at SNR = 30 is shown in Fig. 5a. Here, the red arrow represents the true axon orientation. Overall, SBI (both full and reduced) yields more accurate axon angle estimates than NLLS (tighter circles).

**Fig. 5:**
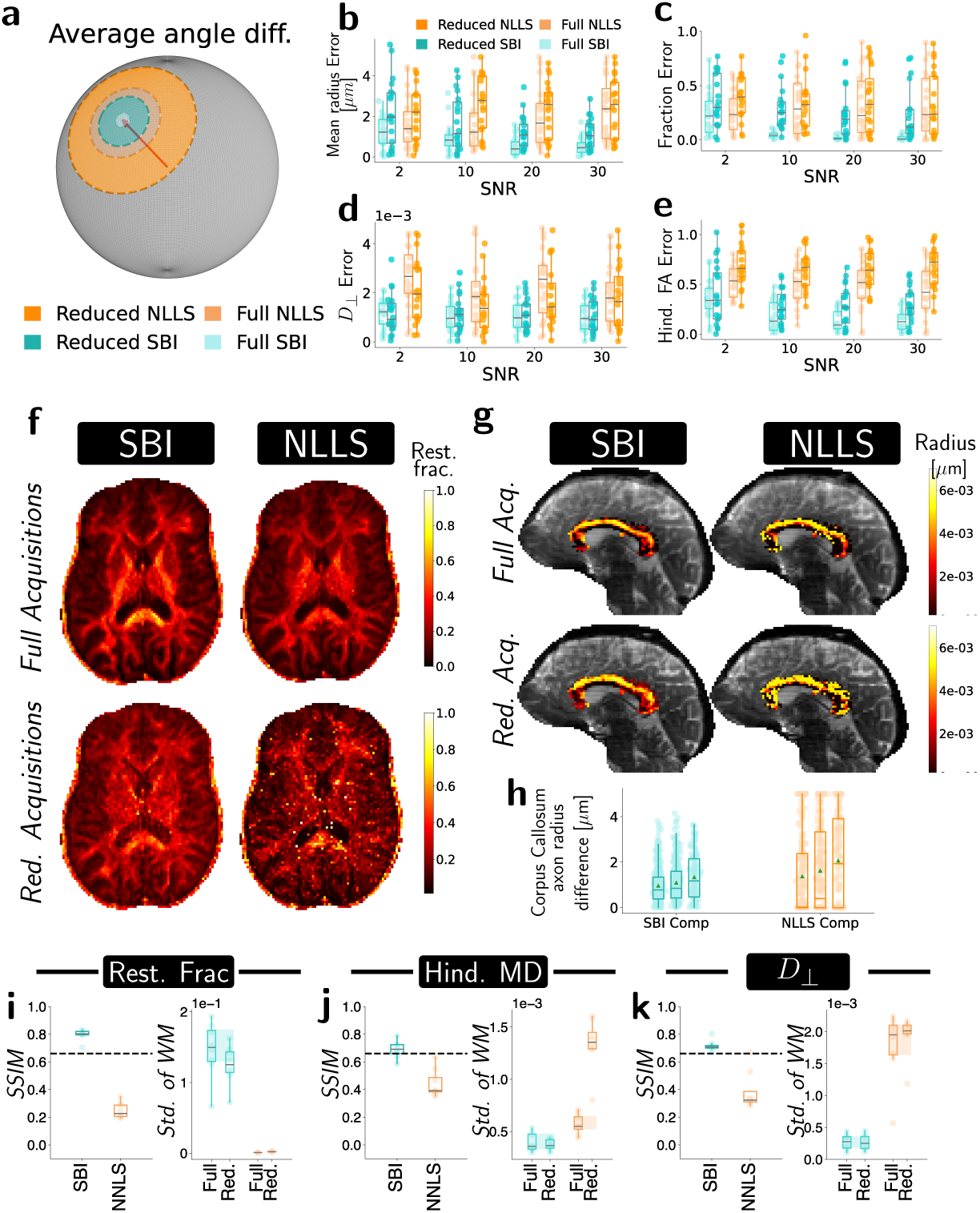
Performance of SBI and NLLS for in-silico and in-vivo AxCaliber data. **a)** For 20 simulated signals with SNR = 30, we calculated the average difference in the angle of the restricted compartment for each fitting routine. The red arrow indicates zero error. Increasing circle sizes denote errors (in radians) of 0.065, 0.24, 0.34, and 0.65 for full SBI, reduced SBI, full NLLS, and reduced NLLS, respectively. **b-e)** Absolute differences between fitted and true parameters for (b) mean radius, (c) restricted fraction, (d) perpendicular diffusion, and (e) hindered fractional anisotropy, using SBI (teal) and NLLS (orange). Black lines show medians; boxes, IQR; whiskers, 1.5 IQR. **f)** Example restricted signal fraction fits for one axial slice, comparing SBI and NLLS. **g)** Example coronal slice of the same individual, with corpus callosum–average axonal radius overlay.**h)** Similarly, segmenting the corpus callosum in two more individuals permits quantitative comparison of reduced versus full acquisition sets (ground truth). Green triangles indicate means; middle lines, medians; boxes, IQR; whiskers, 1.5 IQR. **i)-k)** Structural Similarity Index of (i) restricted fraction, (l) Hindered Mean diffusivity, and (l) rest. perpendicular diffusion, comparing reduced to full acquisitions for SBI (teal) and NLLS (orange).

We then extended our analysis to assess the effects of varying SNR levels on parameter estimation. Signals were generated and corrupted at SNR levels of 2, 10, 20, and 30. For each noise level, we computed the absolute differences between the true and fitted parameters, focusing on the mean radius, restricted signal fraction, perpendicular restricted diffusion, and hindered FA (Figs. 5b–e). In most cases, SBI substantially outperformed NLLS. As expected, increased SNR generally led to reduced errors, indicating improved accuracy with lower noise. However, while full SBI consistently yielded the best performance, NLLS errors did not decrease in the same way. This confirms that for certain parameters — such as mean radius, whose contribution may be small relative to the noise — NLLS struggles to produce accurate estimates, even at SNR = 30.

We next applied the AxCaliber framework to the real-world data. Examples of parameter fits for one individual are shown in Fig. 5f (restricted fraction) and Fig. 5g (average axon radius). Both methods yield similar restricted fraction maps under the full acquisition setting. With reduced data, however, SBI retains more structural detail and exhibits lower noise levels than NLLS.

As illustrated in Fig. 5g, SBI successfully captures the known structure of the axon sizes of the corpus callosum — revealing larger fibres in the body relative to the genu and splenium— for both the full and reduced acquisition sets. Although the reduced acquisition introduces some noise in the axon size fit, it consistently recapitulates the overall structure with the full set and previous findings [58]. In contrast, NLLS shows substantially more noise in its reconstruction and less detailed structure (both full and reduced sets).

We repeated this analysis for the other two healthy individuals and quantified the absolute differences between the inferred axon radius in the corpus callosum and the ground truth (defined by their respective full fit). The results, depicted in Fig. 5h, show that SBI reduced acquisition fits (teal) yield both smaller average and range of errors than NLLS (orange). Importantly, the NLLS fits show a large amount of voxels with the maximum possible error (clustering around 5 *µm*) implying that the reduced NLLS is unable to consistently obtain the correct axon size.

We next also evaluated SBI on additional key AxCaliber metrics for the full set of 8 individuals to demonstrate that the framework works both for healthy and pathological settings. For this, an axial slice from each individual was selected, and the SSIM computed as a proxy for image accuracy. The results (Figs. 5h–k) compare the reduced acquisition results to their full parameter counterparts. We note that SBI outperforms NLLS across all metrics. Particularly in the SSIM of the restricted fraction, while NLLS breaks down, SBI achieves average SSIM values above 0.8, implying a strong resemblance in the results. Given that all the other average SSIM values do not go below 0.7, even with only 10% of the available information for the fit, substantial structural insight is preserved.

## 4 Discussion

In this work, we evaluated how SBI, specifically NPE, performs across mathematical and biophysical models (DTI, DKI, and CHARMED/AxCaliber) under significant noise and limited acquisitions. We trained neural networks on large simulated datasets derived from the mathematical formulations of these models. This approach enhances parameter estimation accuracy while minimizing acquisition time, offering a robust and flexible tool for diffusion MRI analysis. Importantly, our results have the potential of solving a long-standing issue in the field: the huge gap between researcher’s and clinicians’ MRI, stemming from the fact that advanced methods are too time-consuming to be included in the clinical practice. As a consequence, clinical diagnosis in the daily practice of radiology services is still carried out with basic approaches. For example, in the current registered clinical trials in EU, only 3 out of more than 4000 using MRI employ an approach that goes beyond DTI [64]. SBI-based fitting can facilitate more widespread adoption of advanced dMRI approaches like DKI or AxCaliber in clinical applications, providing deeper insights than basic protocols typically used. Additionally, this framework enables the recovery of previously acquired data with short acquisition protocols, like those employed in valuable longitudinal initiatives started decades ago, or affected by noise [65]. Critically, all parameter estimates are obtained without relying on real data, avoiding concerns related to data availability and ethics [36, 37, 54, 55]. Our results could facilitate the use of advanced dMRI approaches like DKI or AxCaliber in clinical applications, providing deeper insights than basic protocols typically used. While we demonstrated this work only on one biophysical model, this approach is inherently flexible and readily extendable to emerging or future models [36].

This work fits within a broader trend of applying neural networks to dMRI tasks, including combining Bayesian frameworks, reconstructing data from undersampled acquisitions, fiber tracking, and enhancing resolution [39, 66–69]. Some works [40, 70] considered using fewer acquisitions but i) only considered simpler models like DTI and ii) relied on real data for training, which can introduce issues with data availability, bias, and generalizability. In terms of SBI, one example of this approach is [36] who demonstrated that SBI is effective for parameter estimation in dMRI, providing quantifiable measures of model degeneracy not available with traditional methods. Other studies using NPE include [71] for epilepsy models, [72, 73] for intravoxel incoherent motion (IVIM) models and [74] who used it to accurately estimate white matter fibre orientations.

Our approach distinguishes itself by combining NPE with dMRI to study AxCaliber and DKI, beyond the linear DTI framework, assessing different noise levels and evaluating minimum acquisition protocols without requiring real data for training. This offers a flexible and efficient framework for accurate parameter estimation when acquisition time is limited. Unlike previous NPE approaches that use summary statistics via multi-layer perceptrons (MLPs) [35, 36], we directly input the diffusion signal to the network, eliminating the need for additional network training and reducing complexity. Two caveats arise: i) our neural networks are specific to the acquisition scheme, and ii) the input dimension can be large with full acquisitions. We addressed i) by providing an identifier for the acquisition scheme (via an integer flag), and for ii) noted that even full acquisition sets were computationally feasible on a personal computer. Future work will aim to incorporate summary statistics to retain simplicity and interpretability while gaining generalization over acquisition numbers and directions.

Nonetheless, our approach may have limitations that warrant further investigation. For example, the use of choice of prior may influence the result of our inference. For DTI, we used simple uniform distributions needing no pre-tuning while for DKI, priors initially required knowledge of parameter distributions from one individual. However, we showed these priors generalized across other individuals, datasets and pathological subjects [42], even with different b-values and protocols, suggesting minimal prior knowledge may suffice. Importantly, in the AxCaliber framework, using pre-reported values from existing literature for the prior distributions of the biological parameters proved sufficient to obtain robust model estimates. Additionally, we acknowledge that NLLS may be a weak baseline to compare SBI to - as its sensitivity to noise is a well-known problem. While NLLS remains one of the most widely used estimators in major toolboxes (for example, DIPY) and clinical studies because of its simplicity, interpretability, and low computational cost, it is limited by noise sensitivity and lack of regularization. Our goal with SBI was to match NLLS’s efficiency while improving robustness. However, we further validated our work by benchmarking the efficacy of SBI in reconstructing kurtosis markers (arguably the most unstable due to non-linearities in their formulation) against three other methods: weighted least squares, a Rician-unbiased approach, and a variational Bayesian approach.

In summary, this work demonstrates how data-friendly SBI methods can substantially minimize acquisition times in both research and clinical settings. By reducing acquisition times by up to 90% in DTI, 85% in DKI and 93% in AxCaliber while maintaining high accuracy, SBI addresses the key limitation of long scan durations in MRI. SBI’s robustness to noise and effectiveness with fewer data points make it especially valuable where high-quality data acquisition is challenging. Demonstrating efficacy in DTI, DKI and AxCaliber, complex models that are more sensitive to noise, opens avenues for impactful clinical research and real-world deployment in dMRI that, after more than 30 years since its first introduction, is still lacking.

## Supporting information

All supplemental figures

## Data and code availability

All data and code is available in the github repository: https://github.com/TIB-Lab/SBIDTI. Within that repository we also provide a link to the trained networks that were used in this work. The MS Data used in the preparation of this work were obtained at the Martinos Center for Biomedical Imaging at Massachusetts General Hospital (MGH) for the NIH 1R21NS123419 Project (Principal Investigator: Caterina Mainero, M.D., Ph.D.). Full MS data can be found at https://digital.csic.es/handle/10261/340684.

## Acknowledgements

We acknowledge the support of the Institute of Neuroscience, CSIC-UMH. We thank Pedro Gonçalves and Caterina Mainero for fruitful discussions and guidance. This research was funded by ”la Caixa” Foundation with Project-ID LCF/BQ/PI23/11970039 (MFE). SDS was supported by the the Spanish Ministerio de Ciencia e Innovacíon, Agencia Estatal de Investigacíon (PID2021-128909NA-I00 and CNS2023-14488), by the Programs for Centres of Excellence in R&D Severo Ochoa (CEX2021-001165-S), by the Generalitat Valenciana through a Subvencion para la contratacíon de investigadoras e investigadores doctores de excelencia 2021 (CIDEGENT/2021/015), and by the Fundacion Pasqual Maragall (ref. 2023-1296).

## Author Contributions

Conceptualization - MFE, SDS. Data curation - MFE. Formal analysis - MFE. Funding acquisition - MFE, SDS. Investigation - MFE, SDS. Methodology - MFE. Project administration - SDS. Resources - SDS. Software - MFE, SDS. Supervision - SDS. Validation - MFE. Visualization - MFE, SDS. Writing – original draft - MFE, SDS. Writing – review and editing - MFE, SDS.

