## Supplementary material for "Simulation-Based Inference at the Theoretical Limit: Fast, Accurate Microstructural MRI with Minimal diffusion MRI Data": All supplemental figures

### Supplemental material

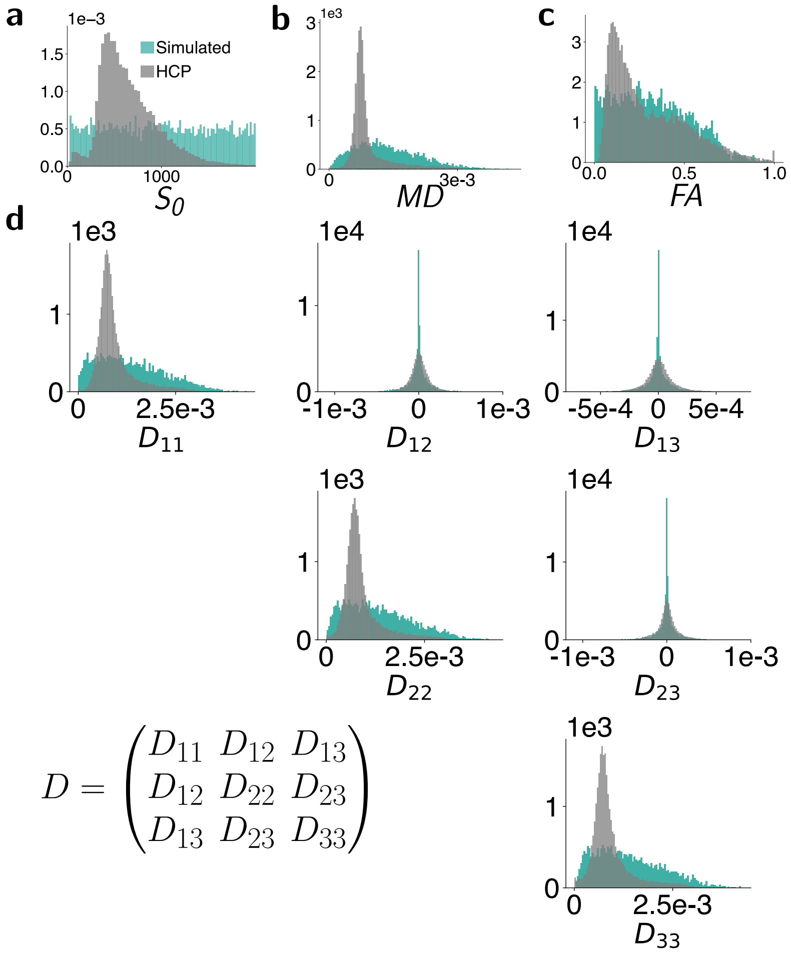


**Fig. S1**: **The distributions of key features in the simulated DTI signals (teal) closely mimic those of the real dataset (gray). a)** Distribution of $S_0$. **b)** Distribution of the mean diffusivity. **c)** Distribution of the fractional anisotropy. **d)** The entries of the simulated diffusion tensor matrix (shown in the bottom left corner), generated as described in the Methods section, span the space of the DTI entries of the HCP data obtained with a NLLS fit.

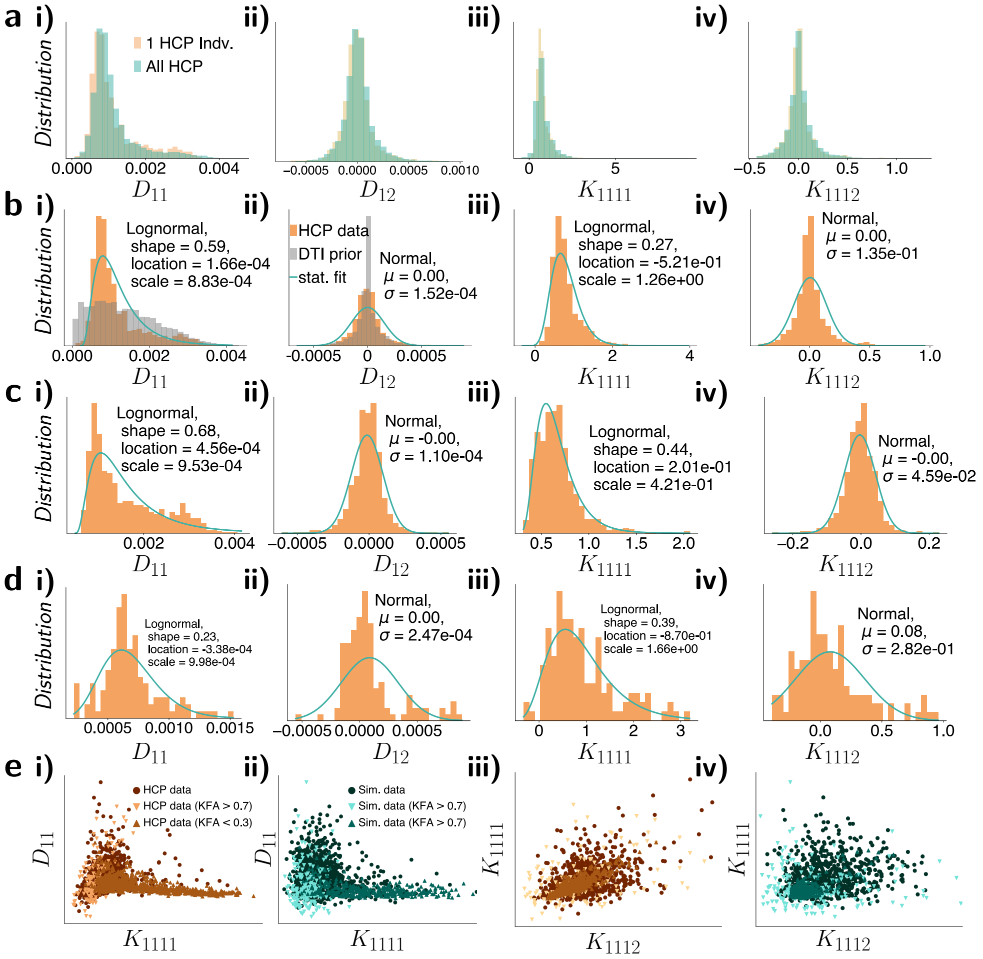


**Fig. S2**: **The DTI and DKI metrics are distributed either normally or log-normally and demonstrate complex non-linear relationships. a)** Distributions of $D_{11}, D_{12}, K_{1111}$ and $K_{1112}$ terms of the single individual (orange) and all individuals (teal) combined from the HCP dataset. **b)** Distribution of the $D_{11}, D_{12}, K_{1111}$ and $K_{1112}$ terms of the single HCP individual (fit with NLLS) in orange. Gray in i) and ii) depicts the results of the simulated DTI values if we sampled from a uniform distribution. Teal line is the best log-normal or normal fit. **c)** Same as b, except that tensors that have KFA >0.3 are filtered out. **d)** Same as b, except that tensors that have KFA <0.7 are filtered out. **e)** Plotting the true $D_{11}$ and the $K_{1111}$ (i), simulated $D_{11}$ and the $K_{1111}$ (ii), true $K_{1111}$ and $K_{1122}$ (iii) and simulated $K_{1111}$ and $K_{1122}$ (iv) entries of the HCP tensors against each reveals a complicated highly non-linear structure, with distinct regions for different KFA values.

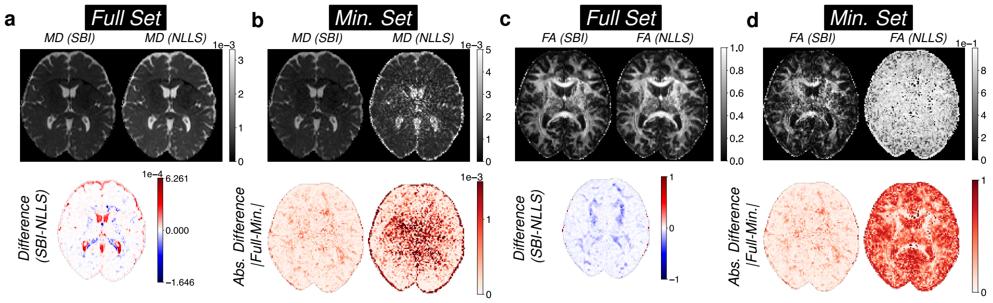


**Fig. S3**: **Using the original voxel resolution of $1.5\mu m$, we demonstrate that SBI's superior performance is independent of resolution. a–d)** Reproduction of the figures in Fig. 2 with the original higher resolution. **c)** Example set of gradient directions of the minimum set of acquisitions. The red arrows represent the random set, i.e. directions were not chosen to maximise electrostatic propulsion. **d)** The resulting MD and FA errors of the minimum set for optimal and random directions (hatched) with increasing SNR compared against the known ground truth. **e)** Comparing the random direction fits against their own optimal direction fits. **f-g)** Fitting MD and FA to one individual of the HCP dataset using the minimum set of acquisitions.

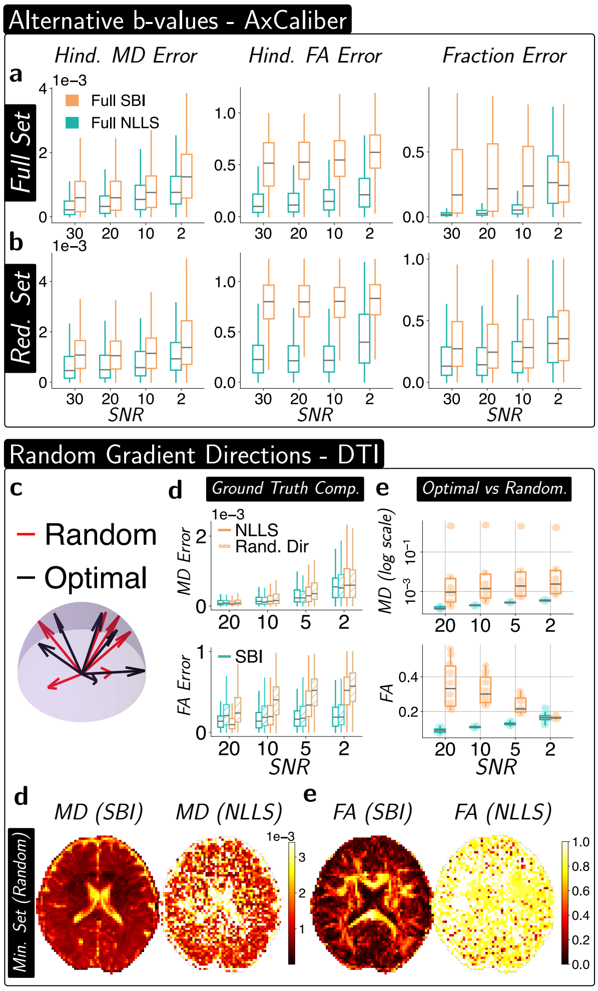


**Fig. S4**: **In-silico analysis of SBI and NLLS fittings of the AxCaliber framework with alternative b-values (b = 3000 and 6000) as well as sub-optimal gradient directions. a-b)** By simulating 500 signals using the AxCaliber framework with the two new b-values, we are able to demonstrate that the resulting errors (here we show a subset of metrics) of SBI outperform the NLLS fits both in the full and reduced set of acquisitions. Box plots represent median values, with boxes showing the IQR and whiskers extending to 1.5 times the IQR.

### Additional DTI errors:

Given a true estimate of the diffusion tensor $D_{\text{true}}$ and a guess for the diffusion tensor $D_{\text{guess}}$, additional tensor specific errors in the DTI setting were calculated as follows:

- Eigenvalue Error= $\sqrt{\sum_{i=1}^{3}(\lambda_{\text{true},i} - \lambda_{\text{guess},i})^2}$, where the eigenvalues of the true and guess tensor were sorted according to their magnitude.
- Tensor error = $\sqrt{\sum_{i=1}^{3}\sum_{j=1}^{3}(D_{\text{true}} - D_{\text{guess}})^2_{i,j}}$

The signal specific errors are, after reconstructing the guess signal $S_{\text{guess}}$ to recreate the true signal $S_{\text{true}}$ given the noisy signal $S_{\text{noisy}}$, defined as:

- $X$ signal Error = $\sqrt{(S_{\text{guess}} - S_{\text{X}})^2}/N$, where $X$ refers to the noisy or true signal and $N$ is the number of acquisitions for the fitting.
- $X$ signal corr. is the Pearson correlation coefficient between the guess signal and the noisy or true signal.

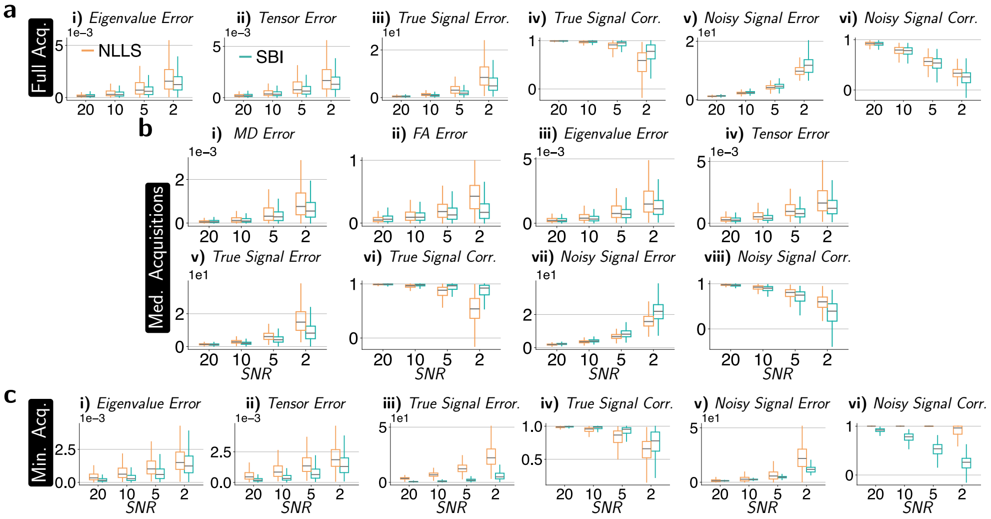


**Fig. S5**: **Similarly to Figure 2, we can perform NLLS and SBI fits using only the medium set of acquisitions. Additional metrics provide insights into the decrease in performance of NLLS versus SBI for the full and minimum set of acquisitions. a)** Using the full set of acquisitions, both NLLS and SBI are fitted to 500 simulated signals. The resulting DTI tensors are compared across six metrics: *i)* difference in sorted eigenvalues, *ii)* difference in Frobenius norm of tensors, *iii)* integrated difference between true (noiseless) and reconstructed signals, *iv)* Pearson correlation between reconstructed and true signals, *v)* integrated difference between noisy and reconstructed signals, and *vi)* Pearson correlation between reconstructed and noisy signals. Box plots represent median values, with boxes showing IQR and whiskers extending to 1.5 times the IQR. Orange denotes NLLS values; teal denotes SBI values. **b)** Repeating this analysis using the medium set of acquisitions and evaluating all metrics (cf. Fig. 2 and a). Box-plots as in a. **c)** Similar analysis as in a, but using the minimum set of acquisitions.

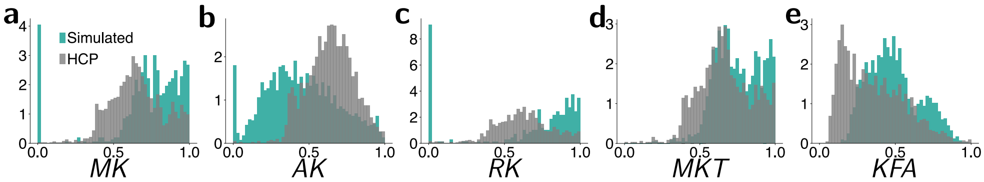


**Fig. S6**: **The distributions of key features in the simulated DKI (teal) mimic those of the real dataset (gray). a)** Distribution of $S_0$. **b)** Distribution of the mean diffusivity. **c)** Distribution of the fractional anisotropy. **d)** The entries of the simulated diffusion tensor (DT) matrix (shown in the bottom left corner), generated as described in the Methods section, span the space of the DTI entries of the HCP data obtained with a NLLS fit.

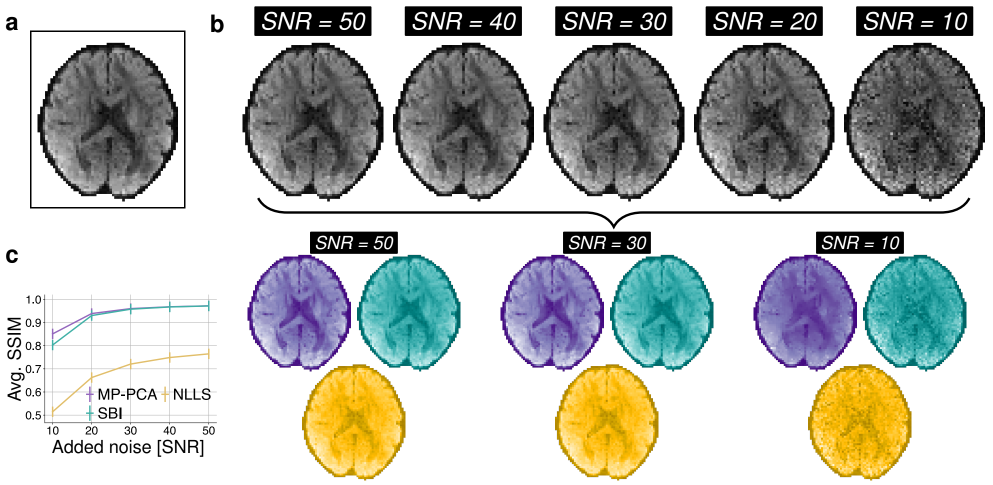


**d**

Fit results

| | | Noise added [SNR] | | | | |
|---|---|---|---|---|---|---|
| | **Metric** | **10** | **20** | **30** | **40** | **50** |
| MD (SBI) | STD | 14e-5 | 7.7e-5 | 5.3e-5 | 4.1e-5 | 3.2e-5 |
| | SSIM | 1.0 | 1.0 | 1.0 | 1.0 | 1.0 |
| MD (NLLS) | STD | 27e-5 | 17e-5 | 13e-5 | 10e-5 | 8.4e-5 |
| | SSIM | 0.95 | 0.96 | 0.96 | 0.96 | 0.96 |
| FA (SBI) | STD | 0.14 | 0.09 | 0.06 | 0.05 | 0.04 |
| | SSIM | 0.40 | 0.60 | 0.67 | 0.70 | 0.71 |
| FA (NLLS) | STD | 0.14 | 0.12 | 0.1 | 0.09 | 0.08 |
| | SSIM | 0.18 | 0.21 | 0.22 | 0.22 | 0.23 |

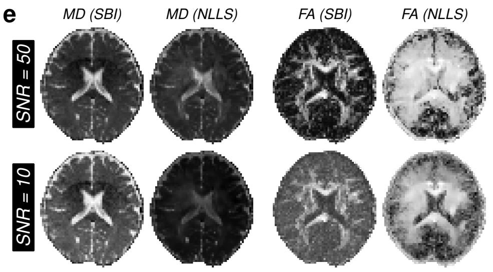


**Fig. S7**: **SBI can also serve as a denoising tool, as long as the model bias is taken into account a)** Example acquisition of one individual from the HCP dataset. **b)** The minimum acquisitions set from this individual were then corrupted with increasing noise (SNR = 10 - 50). **c)** Results of the denoising, using MP-PCA (purple), NLLS (yellow) and SBI (teal) denoising. In the latter two cases, the corrupted signal was fit to a given model (here we use DTI) and then the acquisition reconstructed. SSIM comparison of the uncorrupted and reconstructed acquisitions is also shown in (d). **e)** DTI fit results of (i) SBI directly applied to the corrupted signals and (ii) NLLS applied to the MP-PCA denoised images.

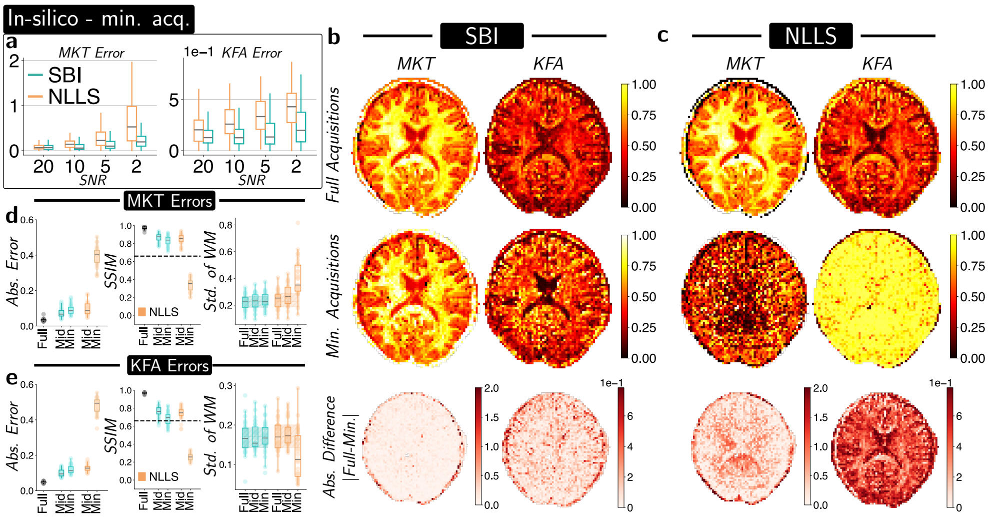


**Fig. S8**: **SBI is able to resolve kurtosis tensor specific metrics even with minimum acquisitions. a)** Mirroring the result of Fig. 2f and h for the MKT and KFA fits. **b)** Applying SBI to the axial slice of an individual (different from the one used for the prior generation) and using the full set of acquisitions provides qualitative comparison of the mean of the kurtosis tensor and the kurtosis fractional anisotropy. For each of the metrics we compute the absolute difference between the full and minimum set. Darker red hues refer to larger differences. **c)** Similar to panels b), but using NLLS. **d-e)** By fitting both SBI and NLLS using different amounts of orientations (Full set, medium and minimum) the medial axial slice of 31 different individuals of the HCP dataset (using one single network), provides insight into the performance of the two methods. Similar to the DTI calculation, the absolute error (i), the SSIM of the axial slices (ii) as well as the standard deviation of the respective metric in the white matter (iii). Given a lack of an objective ground truth, the full NLLS and full SBI results are used as ground truths for their respective subsets. A comparison between the full SBI and full NLLS for the absolute error and SSIM is shown in the gray boxplot. Boxplots represent the median values while the boxes represent the IQR and the whiskers 1.5× the IQR. Orange refers to the fit results the NLLS and teal to the SBI values. The dashed black line in (ii) refers to our acceptable minimum accuracy. The shaded areas in the standard deviation plot refer to the interquartile range of the full fit.

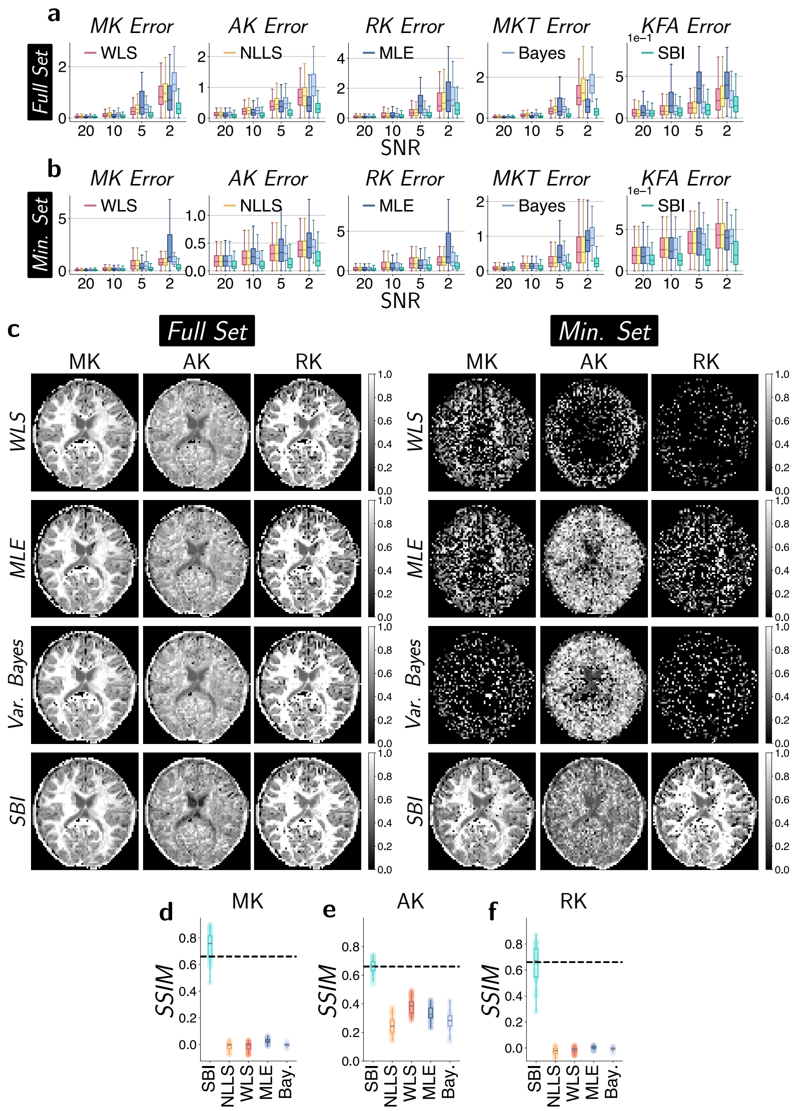


**Fig. S9**: **Further comparison of SBI against other state-of-the-art methods with in-silico and in-vivo data (a–b)** Using the same errors as in Fig. 2c and f, we can evaluate the performance of SBI (teal) against WLS (red), MLE (dark blue) and variational Bayes (light blue). Across a range of SNR values, SBI outperforms these other methods to obtain smaller errors for both the full and minimum acquisition set. Box plots represent median values, with boxes showing the IQR and whiskers extending to 1.5 times the IQR. **(c)** Applying these methods to a real-world HCP slice further demonstrates the robustness of SBI against other more established methods. While all the methods are able to obtain accurate fits when presented the full acquisition set, only SBI provides structure for the minimum acquisition version.**d)-f)** Applying these methods to the 31 HCP subject and evaluating the MK, AK and RK.
